# *Ca. Steroidedax gorgoniicola,* a heterotrophic coral associate with horizontally acquired genes from Endozoicomonadaceae

**DOI:** 10.64898/2026.08.06.743379

**Authors:** Vohsen Samuel, Herrera Santiago

## Abstract

Corals associate with many bacteria whose evolutionary histories and holobiont roles are unknown due to a lack of genomic resources. An example is the BD1-7 clade, which is found in some microbial metabarcoding libraries of corals and has been speculated to be phototrophic. To evaluate its phylogenetic position and assess its metabolic capabilities, we assembled and annotated the genome of an octocoral associate classified as BD1-7. Its full genome revealed that it instead represents a distinct and divergent clade of widespread coral associates. We propose the name *Ca. Steroidedax gorgoniicola* for this associate of *Swiftia exserta*. Unlike the true BD1-7 clade, its genome encoded no pathways to generate ATP from light and instead reveals that it is likely a heterotroph that can degrade steroids, chitin, and collagen as well as produce toxins or antimicrobial compounds and detoxify several reactive oxygen and nitrogen species. In addition, we identified several genes that were likely horizontally transmitted from Endozoicomonadaceae, including transposases and genes involved in virulence and cell adhesion. This work sheds light on the potential role of horizontal gene transfer in the evolution of symbiosis and highlights the importance of obtaining genomes to resolve coral-associated lineages and their metabolic capabilities.

## Introduction

Corals associate with an impressive diversity of microbes [1–3]. Deciphering the roles and evolutionary histories of these associates will advance a more comprehensive understanding of coral ecology as well as aid conservation efforts. For example, studies have shown that diazotrophs can contribute a significant amount of nitrogen to the coral holobiont [4–6], bacterial communities can influence their host’s bleaching tolerance [7], and microbial introductions have occurred that may pose a threat to ecosystem stability [8, 9]. Despite these findings, the roles and evolutionary histories of the majority of coral-associated microbes remain unknown, but these gaps in knowledge may be addressed by generating genomic resources. For example, several genomes have been sequenced of members of the family Endozoicomonadaceae that were isolated from corals [10–14]. Endozoicomonadaceae is one of the most common coral associates, dominatingthe microbiome of many species, including stony corals, black corals, and octocorals, as well as many other marine invertebrates [15]. Obtaining these genomes has accelerated research on Endozoicomonadaceae in several ways by identifying that they likely degrade chitin [16] and produce the protective compound DMS [14, 17], revealing multiple lineages that differ in metabolic capabilities [10, 18], and allowing experiments to identify their physiological changes when initiating symbiosis [19].

Advances such as these are lacking for most coral-associated bacteria, especially those associated with corals in deeper water such as the mesophotic zone. An example is another gammaproteobacterial lineage, the BD1-7 clade. The BD1-7 clade has been reported as a dominant member in the microbiomes of some mesophotic octocorals [20–22] and stony corals [23–25] from the Mediterranean, Red Sea, and South China Sea. At nearby locations, other studies have detected BD1-7 in corals at low abundance [26–28]. Beyond corals, the BD1-7 clade has also been reported in sponges [29], squid gills [30, 31], serpulid worms [32], in association with algae [33–36], and free-living in marine sediment and seawater [37, 38].

Recently, the putative BD1-7 clade was found to dominate (up to 99% 16S relative abundance) the microbiomes of a mesophotic octocoral from the Gulf of Mexico, *Swiftia exserta* [39]. *Swiftia exserta* is an ecologically important species that is the focus of extensive scientific research. It is a major habitat-forming species in the mesophoticzone throughout its range, also occurring from northern Brazil to the mid-Atlantic coast [40, 41]. In the Gulf, it was impacted by the Deepwater Horizon oil spill and is a target of restoration efforts [41, 42]. It is also used as a model system to study the immune response of cnidarians [43–46]. Since coral symbionts have been shown to influence host health and function, it is important to decipher the role of the dominant associates of this important coral species.

In animal hosts, the function and evolutionary history of the BD1-7 clade are unknown. Originally, the BD1-7 clade was named after a novel 16S rRNA sequence obtained from a deep-sea sediment sample from 1,159 m deep [37]. Then, a close relative (HTCC2143) was isolated from seawater (10 m deep) using methods to exclusively cultivate oligotrophic bacteria using low-nutrient media [38]. Cultivation demonstrated that HTCC2143 is psychrophilic or mesophilic and probably obligately oligotrophic [38]. In sponges, isolates classified as BD1-7 clade were identified as cholesterol degraders [29]. In corals, some have speculated that they are phototrophic because the genome of HTCC2143 encodes proteorhodopsin [22, 23, 47]. Consistent with this speculation, the dominant ASV classified as BD1-7 clade strongly decreased in relative abundance with depth in *Swiftia exserta* [39].

In order to assess the metabolic capabilities and evolutionary history of the putative BD1-7 associate found in *Swiftia exserta,* we sequenced metagenomes, conducted phylogenomic and comparative genomics analyses, and screened metabarcoding datasets of corals for relatives to understand the ecological breadth of this lineage.

## Results

### Genome Statistics and Phylogenetics

We successfully assembled a 4.3 Mbp genome for the bacterial associate of *Swiftia exserta* previously classified as the BD1-7 clade (98% completeness, **Table 1**). Phylogenomic analysis based on the bac120 gene set did not place this associate of *Swiftia exserta* within the original BD1-7 clade (**Fig. 1A**). Instead, it formed a novel clade with associates from another octocoral (*Leptogorgia sarmentosa*) and sponges, andwas sister to the Sinobacteriaceae. This new clade shared an amino acid identity >60% (**Table S1**), consistent with the level of a family within the Gammaproteobacteria [48]. This was consistent with phylogenetic analysis using the full-length 16S rRNA gene which identified other close relatives from octocorals (*Eunicella cavolini, Gorgonia ventalina,* an unknown bamboo coral), hexacorals (*Tubastrea coccinea* and an unidentified deep-sea black coral), a hydrocoral (*Stylaster aurantiaca*), seawater from hydrothermal vents, and a surgeonfish gut (**Fig. 1B**). The general topology of these trees was consistent with other phylogenetic analyses of gammaproteobacteria including those targeting BD1 -7 clade and Spongiibacteraceae [48–50]. We propose the name *Ca. Steroidedax gorgoniicola* gen. nov. sp. nov. for this associate of *Swiftia exserta* (see below for species description).

**Figure 1.**
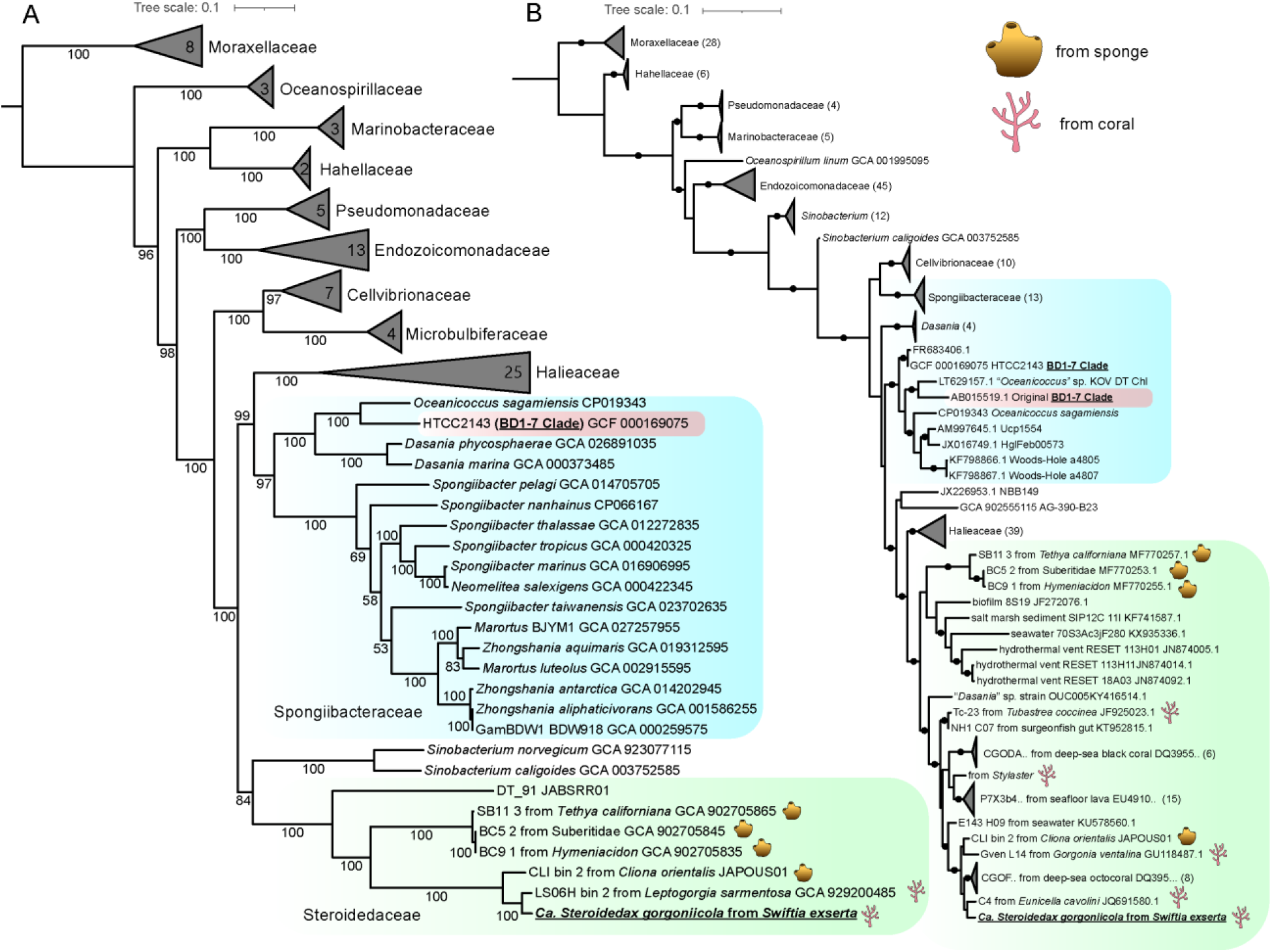
Maximum likelihood phylogenetic trees showing the relative phylogenetic positions of the BD1-7 Clade and the associate of *Swiftia exserta* using (A) an amino acid alignment of the bac120 gene set (B) the 16S rRNA sequences. Numbers at nodes represent Ultrafast bootstrap support values. Black circles in (B) denote nodes with greater than 90%. Numbers in collapsed clades and in parentheses denote the number of genomes or sequences within.

**Table 1.** Genomic summary of *Swiftia exserta’*s dominant associates.

|  |  | *Ca. Steroidedax gorgoniicola* | *Ca. Nereiplasma swiftiae* |
| --- | --- | --- | --- |
| CheckM | Size | 4,273,284 bp | 686,998 bp |
|  | GC content | 40.8% | 22.0% |
|  | Longest scaffold | 110,275 | 73,867 bp |
|  | N50 | 23,592 bp | 35,875 bp |
|  | # scaffolds | 348 | 45 |
|  | Predicted genes | 3,740 | 644 |
|  | Coding density | 0.84 | 0.89 |
|  | Completeness | 98.39 | 94.92 |
|  | Contamination | 1.70 | 0.00 |
| DRAM | Predicted genes | 3,628 | 616 |
|  | tRNAs | 41 | 20 |
|  | rRNAs | 3 (5S, 16S, 23S) | 2 (16S, 23S) |

### Distribution in other corals

Using the full 16S rRNA gene of *Ca. Steroidedax gorgoniicola,* we found twenty-two closely related ASVs in several other coral species at high relative abundances (at least 10% in one sample, **Fig. 2A, Table S2**) using a modified microbial metabarcoding database of global cnidarians [1]. These included seven other octocorals (*Scleronephthya gracillima*, *Eunicella cavolini, E. verrucosa, E. singularis, Leptogorgia sarmentosa, Melithaea rubrinodis*, and *Pacifigorgia cairnsi*), six stony corals (*Montipora aequituberculata*, *Merulina ampliata, Orbicella faveolata, Acropora hemprichii, Acropora muricata,* and *Desmophyllum pertusum* [formerly *Lophelia pertusa*]), a black coral (*Antipathella subpinnata*), and a hydrocoral (*Millepora*). These corals were globally distributed (**Fig. 2B**) and the BD1-7 clade was not reported in at least 10 of these coral species. There was little to no overlap across species except across *Eunicella* spp. and *Leptogorgia sarmentosa* which all hosted the same ASV. For every ASV, the phylogenetic position with the highest likelihood was near a member of Steroidedaceae isolated from an animal host and never closest to a free-living isolate (**Fig 2C**).

**Figure 2.**
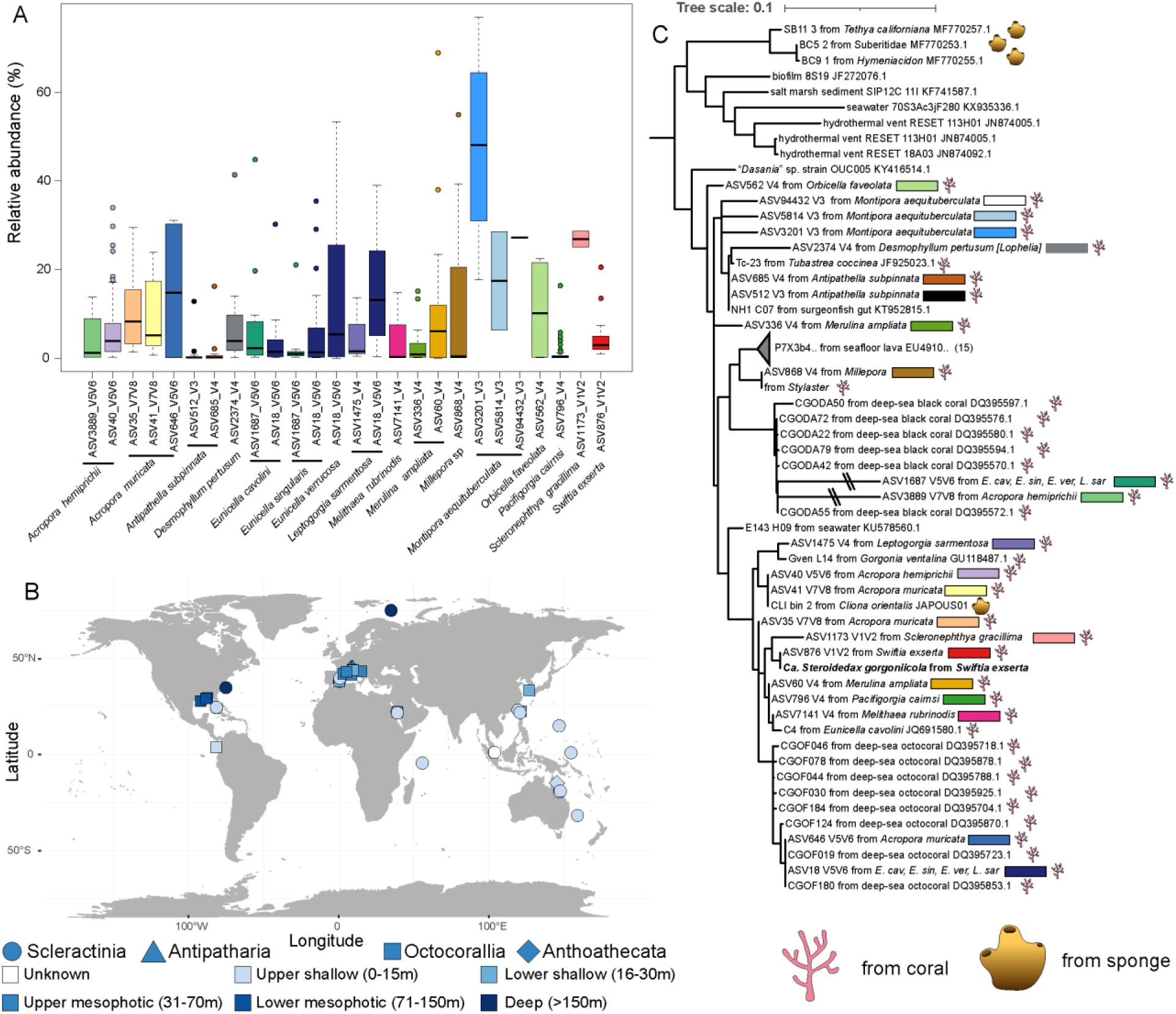
(A) The relative abundances of Steroidedaceae ASVs among corals and (B) the geographic and depth distributions of those samples. (C) The most likely position of each Steroidedaceae ASV within the 16S rRNA reference tree.

### Metabolic characterization of Ca. Steroidedax gorgoniicola

Genome annotation suggests that *Ca. Steroidedax gorgoniicola* is an oxidative heterotroph. Its genome encodes glycolysis, the Pentose Phosphate pathway, the Entner-Doudoroff pathway, and the TCA cycle, suggesting it generates ATP through the catabolism of sugars and fatty acids (**Fig. 3A, Table S3**). It also encoded chitinase and collagenase as well as genes that degrade steroids (cholesterol, testosterone, and other androgens) to generate ATP. Its genome did not encode any pathway or genes suggesting that it could utilize light to generate ATP. It did not contain a rhodopsin (K04643) like HTCC2143. However, it did encode a protein with a BLUF domain, which may sense blue light but is unrelated to rhodopsin [51].

**Figure 3.**
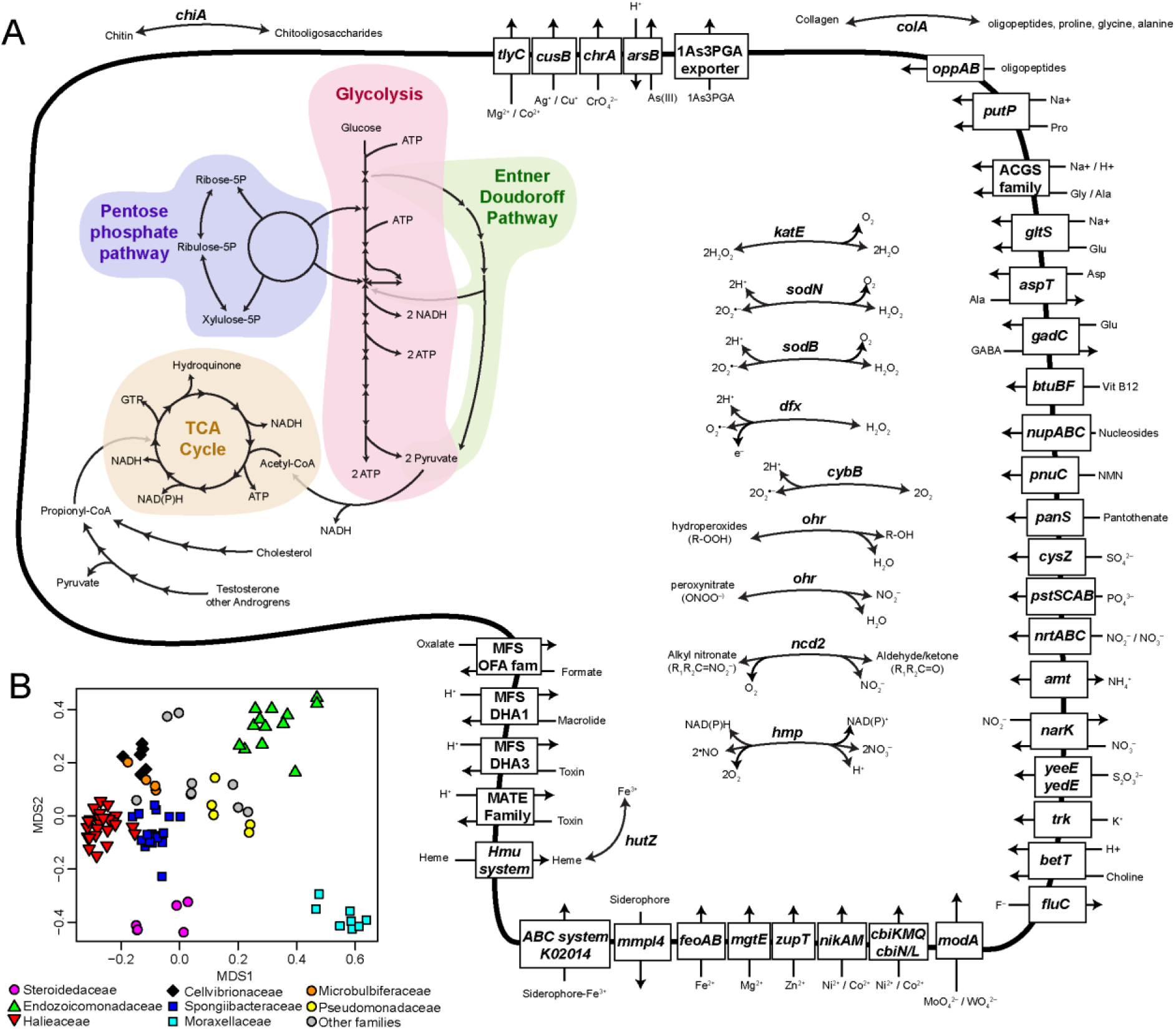
(A) A metabolic map of *Ca. Steroidedax gorgoniicola* highlighting select pathways inferred from genome annotations. (B) A non-metric Multidimensional Scaling plot the gene content of all genomes used in the phylogenomic analysis using Jaccard’s distance of orthogroups present in more than two genomes.

In addition, the genome encoded many genes that detoxify reactive oxygen and nitrogen species (two catalases, two superoxide dismutases, superoxide reductase, two superoxide oxidases, two nitronate monooxygenases, nitric oxide dioxygenase, glutathione peroxidase, Group II and III truncated hemoglobins, lipoyl dependent peroxiredoxin (Ohr), paraquat-inducible proteins (pqiABC) [52–56] as well as many genes to synthesize potential toxins or antibiotics (polyketide cyclases, non-ribosomal peptide synthases, toxoflavin synthases, Rhs family proteins, and an antibiotic biosynthesis monooxygenase). It also encoded genes involved in antibioticresistance(class C beta-lactamases) as well as assimilatory nitrate and sulfate reduction.

Its genome displayed no evidence of reduction, and it encoded the pathways to synthesize all amino acids as well as many cofactors and vitamins: molybdopterin, coenzymes A and F420, vitamins B1, B2, B5, B6, B7, B9, and B12 (thiamin, riboflavin, pantothenate, pyridoxine, biotin, folate, and cobalamin), chorismate, and heme. Other notable genes included agmatine deiminase and spermidine synthase.

### Comparative Genomics of Steroidedaceae

*Ca. Steroidedax gorgoniicola* and its closest relatives were similar to each other and differed from their closest related families in the presence of orthogroups (**Fig. 3B, Tables S3-S4**). Among the 1,279 orthogroups shared by all Steroidedaceae were genes to degrade steroids as well as several genes to protect against reactive oxygen and nitrogen species (superoxide dismutase, superoxide reductase, superoxide oxidase, nitronate monooxygenase, nitric oxide dioxygenase, and truncated hemoglobin). Other notable genes included hemolysins, a vitamin B12 importer, and nitrite reductase. 19 orthogroups were unique to Steroidedaceae and absent from all other genomes used.

### Horizontal gene transfer from Endozoicomonadaceae

Several genes of *Ca. Steroidedax gorgoniicola* showed evidence of horizontal gene transfer from *Endozoicomonadaceae*. We identified 124 genes in which two or more matches to Endozoicomonadaceae genes were among the top ten matches using blastp on the NCBI nr v4 database (**Table S3**). These HGT candidates were closer on average to transposases (Wilcoxon rank sum test p=0.006, **Fig. 4A**) and more spatially clustered than other genes in the genome (p<0.0001, **Fig. 4B**). Gene trees confirmed that most of these (>85, 69%) clustered among genes from Endozoicomonadaceae rather than with genes from the closest relatives of Steroidedaceae (**Table S3**). These genes were not likely to be from mis-binned contigs from co-occurring *Endozoicomonadaceae* since they were on contigs that also included non-HGT candidates (**Fig. 4C**) and did not differ in depth of coverage from other contigs (**Fig. S1**). These genes seem to result from at least three potential HGT events: 1) from *Neoendozoicomonas* spp. to *Ca. Steroidedax gorgoniicola* and/or to its common ancestor with the associate from *Leptogorgia* [43 genes], 2) from an unknown Endozoicomonadaceae lineage to the common ancestor of the Steroidedaceae from *Swiftia, Leptogorgia*, and the sponge *Cliona orientalis* [19 genes], and 3) an older transfer between basal lineages of Endozoicomonadaceae and Steroidedaceae [9 genes] (**Fig 4DE**).

**Figure 4.**
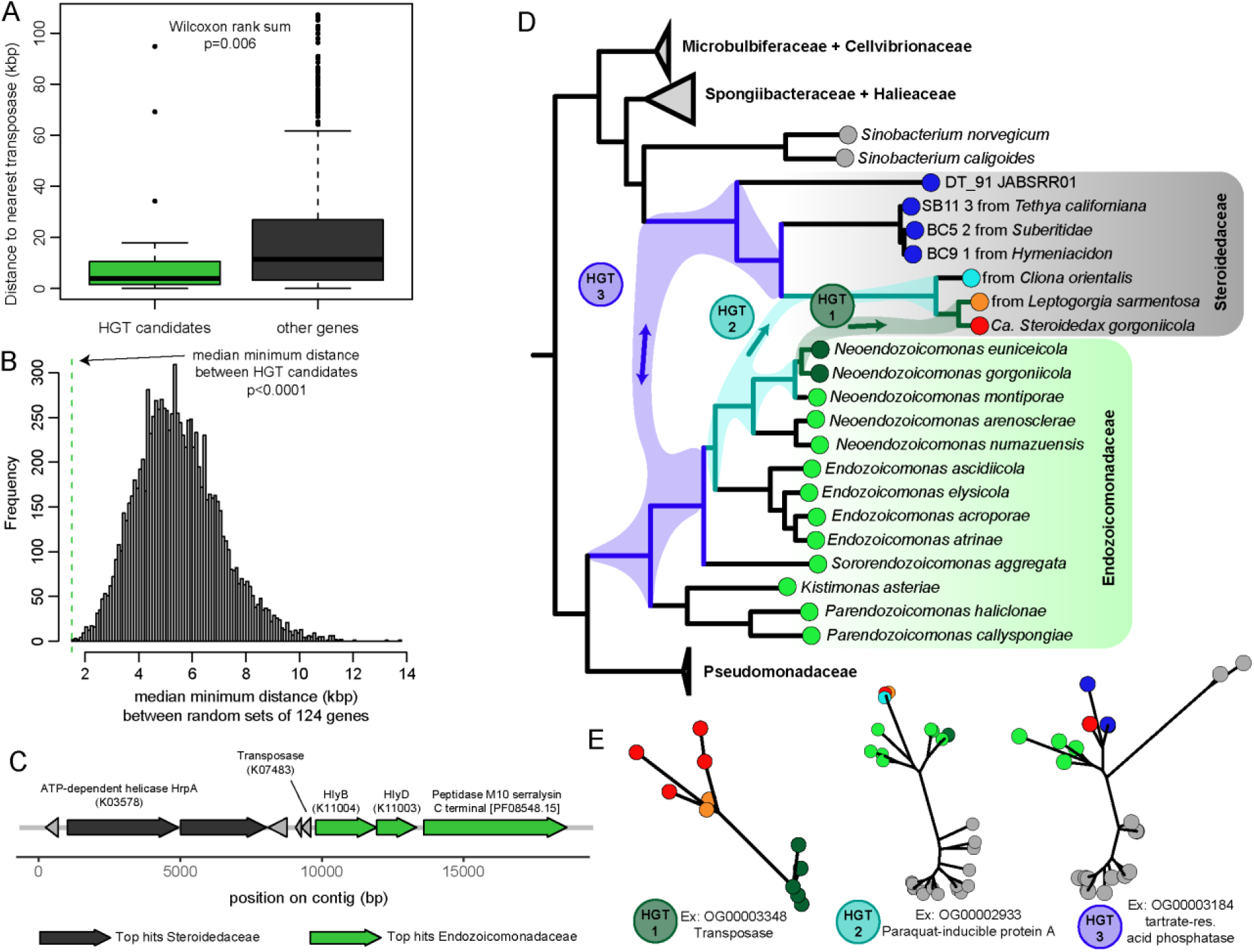
(A) The distribution of distances of HGT candidates to the nearest transposase compared to other genes. (B) The median minimum distance between HGT candidate genes (measure of clustering) compared to the distribution when selecting an equal number of random genes. (C) Example contig with HGT candidates near transposase. (D) Phylogenetic tree demonstrating the direction of the three hypothesized HGT events (E) Example gene trees of orthogroups for each predicted HGT event.

Genes from all three of these groups had annotations involved in defense against the acid, nitrosative, and oxidative stresses characteristic of immune responses: (1) Ohr, (2) pqiA, (3) truncated hemoglobin, and tartrate-resistant acid phosphatase type 5 [57]. Additionally, the most recent event (1) included genes that are typically involved in the mechanism of HGT (transposases, reverse transcriptase, phage tail tube protein, and a gene with DUF4372) as well as genes involved in toxin-antitoxin systems (DUF2442, DUF4160), manipulating hosts or virulence factors (PI-PLC, hemolysin-type type-1 secretion system with an unknown gene encoding the C terminal domain of an M10 peptidase instead of an associated hemolysin, SET domain protein, glycosyl hydrolase family 99), an OprD porin, and an antibiotic biosynthesis monooxygenase [58–61]. Event 2 included the virulence factor, phospholipase C, and genes involved with surface protein synthesis, export, and modification (capsule polysaccharide synthesis and export genes, mannose-1-phosphate guanylyltransferase / mannose-6-phosphate isomerase, UDP-N-acetylglucosamine 2-epimerase, and a giant protein). This giant protein was 11,389 aa long with 40 domain annotations including a peptidase (myxosortase dependent M36 family metallopeptidase) and many domains involved with cell adhesion and host invasion (autotransporter adhesin AidA, 3x Pentraxin, 7x Concanavalin A-like/glucanase superfamily, 7x VCBS repeat, 3x tandem-95 repeat, 7x cadherin, 7x cadherin tandem repeat, 3x FG-GAP repeat / integrin, F5/8 type C domain / discoidin: CD-Search). Finally, the oldest event (3) included a chitinase and several other genes, including UDP-N-acetylglucosamine 4,6-dehydratase, beta-ketoadipyl-CoA thiolase, Acyl-CoA dehydrogenase FadE17, and a gene with an oxidoreductase FAD-binding domain.

### The other dominant microbe, a mycoplasma

In addition to *Ca. Steroidedax gorgoniicola,* we obtained a genome for the dominant mycoplasma in *Swiftia exserta* that was 687 kbp with an estimated completeness of 95% (**Table 1**). A phylogenomic tree using the bac120 gene set clusteredit in a lineage near Metamycoplasmataceaealongsidemycoplasmas isolated from *Eunicella gazella,* and the same samples of *Leptogorgia sarmentosa* that yielded the closest relative of *Ca. Steroidedax gorgoniicola* (**Fig. 5A**). Its genome reveals that it is a fermentative mycoplasma encoding the genes necessary to generate ATP from glycolysis and the arginine deiminase pathway (**Fig. 5B, Table S5**). Typical of Mollicutes, it lacks the ability to synthesize most amino acids. We propose the name *Ca. Nereiplasma swiftiae* for this mycoplasma from *Swiftia exserta*.

**Figure 5.**
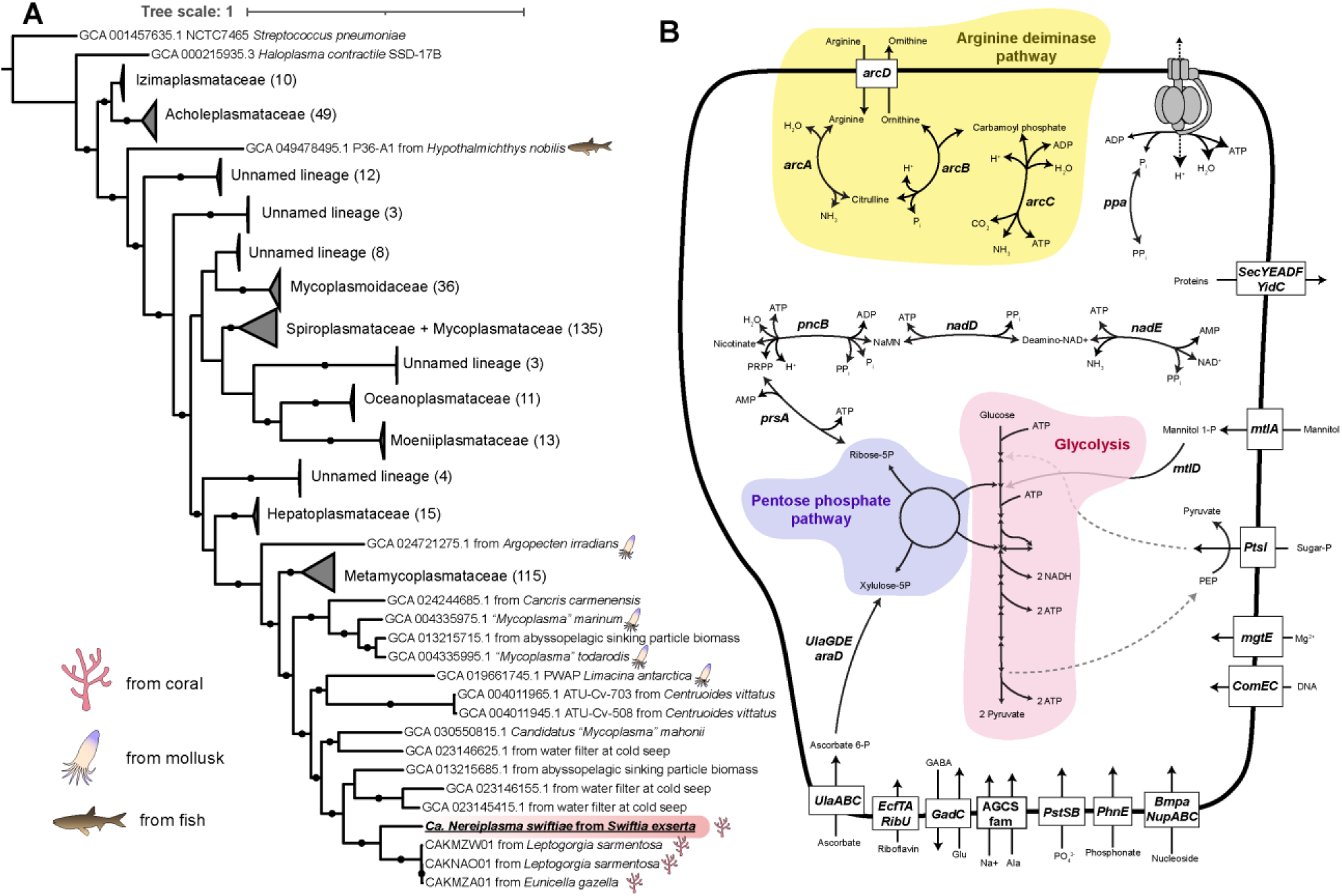
(A) Maximum likelihood phylogenomic tree of *Ca Nereiplasma swiftiae* from *Swiftia exserta* among the Mollicutes using an amino acid alignment of the bac120 gene set. (B) A metabolic map highlighting select pathways inferred from genome annotations.

### Etymologies and species descriptions

#### Steroidedax gen. no

(Ster.oid’.edax), E. n. *Steroid*, a steroid; L. suff. -*edax* devouring, consuming; N.L. neut. n. S*teroidedax*, a steroid-consuming bacterium, referring to its predicted ability to degrade steroids. See description for type taxon: *Ca. Steroidedax gorgoniicola*.

#### Steroidedax gorgoniicola sp. Nov

(gor.go.ni’.i.co.la), N.L. fem. n. *gorgonia*, a gorgonian coral; L. suff. - *cola*, inhabitant, dweller; N.L. n. *gorgoniicola*, inhabiting a gorgonian octocoral. Oxidative heterotroph associate of the gorgonian *Swiftia exserta*. Type material is NCBI Assembly: JCAIKI000000000.

#### Nereiplasma gen. nov

(Ne.re’.i.plas.ma), Gr. masc. n. *Nereus*, a primordial sea god in Greek mythology; Gr. neut. n. *plasma* molded, lacking a cell wall, conventional suffix for mollicutes; N.L. neut. n. *Nereiplasma*, wall-less bacterium of the sea. See description for type taxon: *Ca. Nereiplasma swiftiae*.

#### Nereiplasma swiftiae sp. Nov

(swift.ti.ae’), N.L. gen. fem. n. *swiftiae,* of *Swiftia exserta*. Fermentative mycoplasma associate of the mesophotic octocoral, *Swiftia exserta.* Type material is NCBI Assembly: JCAIKJ000000000.

## Discussion

### Misclassification as BD1-7 clade and prevalence among corals

We found that the dominant associate of *Swiftia exserta* was not closely related to the BD1-7 clade as predicted bythe initial ASV classification [39]. Instead, *Ca. Steroidedax gorgoniicola*, representsa novel family (Ca. Steroidedaceae) that is present in several marine invertebrates, including corals, sponges, annelids, and mollusks. Previously, ASVs classified as the BD1-7 clade had only been reported in a few other coral species. We demonstrated that all of these are actually Steroidedaceae, and we detected them in several more coral species, revealing it is far more widespread than previously understood. In their original publications, these Steroidedaceae ASVs we identified were classified as BD1-7 clade, Gammaproteobacteria, and even *Melitea,* highlighting the limitations of common classification tools applied to novel lineages. In our analysis, the full 16S rRNA gene for *Ca. Steroidedax gorgoniicola* was sufficient to distinguish it as a novel lineage. Incorporating sequences with confirmed phylogenetic backgrounds in classification databases should improve their performance for novel lineages. This should improve the ability to synthesize results across metabarcoding datasets to help identify more lineages that are widespread among marine invertebrates.

### Role of Ca. Steroidedax and interaction with the coral

It was previously speculated that the BD1-7 clade in corals is phototrophic because opsins were encoded in the genome of one BD1-7 clade member. The genome of *Ca. Steroidedax gorgoniicola* did not encode any gene or pathway suggesting it is phototrophic, consistent with the realization that it is unrelated to the BD1-7 clade. Instead, its genome suggests that it is a heterotroph that can degrade steroids, chitin, and collagen. The lack of genome reduction suggests that *Ca. Steroidedax gorgoniicola* has a free-living stage while *Ca. Nereiplasma swiftiae* exhibits the typical genome reduction of Mollicutes and likely relies on its host.

It is still unclear what role *Ca. Steroidedax* may play in the coral holobiont. It is possible that it may play a digestive role by degrading the chitin and collagen from its host coral’s diet of zooplankton or could provide its host with essential compounds. The many antimicrobial compounds it is predicted to synthesize may exclude other microbes from infecting *Swiftia*, including pathogens, and probably underlie its dominance in *Swiftia’s* microbiome. Several of these compounds target bacterial cell walls, which is consistent with the fact that the other dominant microbe in *Swiftia exserta* is the cell wall-less *Ca. Nereiplasma swiftiae*.

It is also possible that *Ca. Steroidedax gorgoniicola* is not a mutualist but is rather a parasite. None of the *Swiftia* sampled in this study nor in Vohsen & Herrera (2024) [39] showed any signs of disease, so *Ca. Steroidedax* does not seem to be a pathogen. Recently, Endozoicomonadaceae has been proposed to potentially play a parasitic role in corals in some contexts [15]. Interestingly, the genome of *Ca. Steroidedax gorgoniicola* possesses many genes shared with Endozoicomonadaceae including chitinase, catalase, genes that synthesize antimicrobial compounds, giant proteins, and genes involved in steroid degradation (shared with *Neoendozoicomonas*). Thus *Ca. Steroidedax* may play a similar role in coral holobionts and may even form large aggregates like Endozoicomonadaceae.

### Horizontal Gene Transfer with Endozoicomonadaceae

We detected horizontal gene transfer from Endozoicomonadaceae into Steroidedaceae in what appears to be at least three events. These include genes typically involved in HGT events, including transposases and viral genes, but also genes critical for successful infection of hosts or virulence, including functions like cell adhesion and defense against the nitrosative and oxidative attacks of the innate immune system.

The most recent HGT event appears to be from *Neoendozoicomonas* spp. isolated from octocorals to an ancestor of *Ca. Steroidedax gorgoniicola* and its closest relative, which is also from an octocoral. These genes include transposases but also several virulence factors including a hemolysin specific type-1 secretion system but instead of encoding a hemolysin, it encoded an unknown gene with the C terminal domain of an M10 peptidase (**Fig. 4C**). Hemolysins are used by pathogenic bacteria to rupture host cells [62] but are also used by nodule forming mutualists to initiate nodule formation in plants and influence host specificity [63–65]. Whereas the family M10 peptidase family includes serralysins and are zinc metalloproteases that degrade host proteins, including the cell-surface adhesion molecules of macrophages to avoid phagocytosis [66, 67]. If this unknown gene or the other virulence factors among the group 1 HGT candidates is specific to octocorals, it is possible that this HGT event allowed the ancestor of *Ca. Steroidedax gorgoniicola* to infect octocorals. This transfer may have occurred while both members were in their free-living stages and was mediated by a virus. Another possibility is that the HGT event occurred when the ancestor of *Ca. Steroidedax* co-infected an octocoral that already hosted *Neoendozoicomonas*. HGT between co-occurring symbionts has been documented in insects, plants, and chemosymbiotic mussels where it is thought to mediate the acquisition of new metabolic capabilities and host specificity [68–71]. But this has not been documented among coral associates. As in these other systems, HGT may allow new microbial lineages to gain the ability to infect corals from lineages already adapted to corals, establishing new symbioses. It may also introduce new metabolic functions to existing coral-associated lineages. We detected virulence genes among all three HGT events, suggesting that HGT from Endozoicomonadaceae may have facilitated the spread and infection of different hosts across most of the evolutionary history of Steroidedaceae.

## Conclusion

Generating genomic resources provides a wealth of critical information regarding coral symbionts. Here, we show it can correct the phylogeny of coral symbionts, support the synthesis of the growing number of metabarcoding datasets by improving amplicon classifications, and resolve questions regarding symbiont capabilities. Further, it can uncover unforeseen aspects of a symbiont’s evolutionary history that could represent important mechanisms in the evolution of symbiosis in corals.

## Methods

### Sample Collection and DNA extraction

Samples were collected as described in Vohsen and Herrera (2024) [39]. Of the samples included in that study, two colonies of *S. exserta* were selected from each of Mountaintop Reef and Geyer Bank where the relative abundances of BD1-7 clade and mycoplasma were high, respectively (**Table S6**). DNA was extracted following a modified salting-out protocol as in Vohsen and Herrera (2024) [39]. To remove PCR inhibitors, DNA extracts were cleaned using a DNeasy Powerclean Pro cleanup kit (Qiagen) with modifications to preserve high molecular weight. Samples were mixed by inversion, instead of vortexing, and DNA was eluted from the spin column twice using 100µL of TE buffer after incubating on the silica membrane for 5 minutes both times. The DNA concentration of the second elution was quantified using a nanodrop and Qubit BR kit on a Qubit 4.0 Fluorometer, then 1µg of DNA was enriched for microbes using an NEBNext Microbiome DNA enrichment kit (New England Biolabs), purified using an ethanol precipitation, and resuspended in 30µL TE buffer following the manufacturer’s guidelines. Microbe-enriched DNA extracts were then sent to Novogene for library preparation and sequencing on an Illumina HiSeq2500 platform using 150bp paired-end sequencing. Additionally, publicly available libraries from five additional *S. exserta* colonies were utilized for metagenome assembly and binning: Alabama Alps Reef (n=2), East Flower Garden Banks (n=2), and Bouma Bank (n=1).

### Metagenomic Analysis

Each library was individually screened using phyloFlash v3.4.1 [72] to obtain an initial assessment of the microbial community and to assemble any SSU rRNA genes. Then, the metagenomic wrapper suite, MetaWRAP [73], was used to assemble metagenome-assembled genomes (MAGs) following its general guidelines. Raw read files were quality filtered with the read_qc module with default parameters. Reads corresponding to coral were removed from *S. exserta* metagenomes using a host genome generated in-house. Reads from four libraries in which phyloflash detected BD1 -7 were pooled and assembled with metaSPAdes v3.13.0 [74]. Following the MetaWRAP pipeline, the remaining unassembled reads were assembled with MEGAHIT v1.1.3 [75], then both assemblies were concatenated. To obtain a mycoplasma genome, the four libraries sequenced here were pooled and assembled as above. The metagenome assemblies were binned using MetaBAT v2.12.1 [76], MaxBin v2.2.6 [77], and CONCOCT [78], refined using the bin_refinement module, and visualized with Blobology [79]. Coral contigs were removed from the BD1-7 MAG with the following rule using the metabat2 coverage output: V1/abs(V2) <1.3 where V1 is the sum of the depths of coverage of a contigacross the fivesamples without BD1-7 and V2 is the average depth of coverage of that contig among the 4 samples with BD1-7 minus the average among the 5 samples without BD1-7. V1 reflects how closely the coverage pattern of a contig matches those likely to be coral while V2 reflects how closely it matches those likely to be from BD1 -7. The threshold separated two clusters of contigs in V1 and V2 space (**Fig. S2-3**) which corresponded to contigs classified as eukaryotic or prokaryotic as well as their associated unclassified contigs with similar coverage patterns. The reassembly module was applied to improve the mycoplasma bin and the strict version was kept. The filtered bins were then annotated using DRAM v1 [80]. Average Amino Acid Identities (AAI) were calculated using EzAAI v1.2.4 [81].

### Phylogenetic Analyses

In order to infer the phylogenetic positions of *Swiftia’s* associates, maximum likelihood phylogenetic trees were constructed using publicly available sequences for assembledgenomes and the 16S SSU rRNA to place our sequences within a larger diversity (**Table S7-8**).For assembled bacterial genomes, GTDB-tk v2.4.0 was used to generate concatenated amino acid alignments of 120 marker genes corresponding to the bac120 geneset, which were used to generate maximum likelihood phylogenetic trees using IQ-TREE v2.3.0 [82] with UFBoot2 [83] bootstrap support values (1000 replicates). Additionally, 16S SSU rRNA gene sequences were aligned using Clustal Omega v1.2.4 [84], and a tree was constructed using IQ-TREE2. To obtain 16S sequences for microbes that are only represented in publicly available MAGs, barrnap v0.9 [85] was used to identify rRNA genes within their assemblies (**Table S8**). Finally, a 16S sequence classified as BD1-7 clade was obtained using phyloflash after pooling publicly available *Stylaster aurantiacus* libraries (**Table S8**).

### Screening the global cnidarian microbial diversity database

We screened each of the five global cnidarian 16S rRNA libraries (V1-V2; V3; V4; V5-6; V7-8) that characterize the microbial diversity of 212 cnidarian species [1] to identify ASVs related to *Ca. Steroidedax gorgoniicola.* Libraries that contained the filtered and denoised sequences before they were trimmed down to a common amplicon length were chosen for this comparison [1]. All ASVs were blasted against the full-length 16S rRNA gene and selected if they matched with at least 94% identity and 95% coverage. These ASVs were then blasted against the NCBI nt v4 database and only retained if they matched a sequence within Steroidedaceae (**Table S9**) with greater than 88% identity, 95% coverage, and 900bp subject length. These ASVs were kept if more than one of their top nine matches based on E-value were within Steroidedaceae and included a match in the top four. These ASVs were then placed in the phylogenetic tree of full-length 16S sequences using clustuneR and pplacer to confirm identity as Steroidedaceae after aligning to the reference sequences using Clustal Omega.

### Comparative genomics and HGT analyses

Clusters of orthologous genes were identified across all genomes used in the phylogenomic analysis above using Orthofinder v2.5.5 [86] after using Prodigal v2.6.3 [87] to extract coding regions. A Non-metric Multidimensional Scaling plot was constructed using Jaccard distances of COGs that were present in more than two genomes through the vegan package in R.

*Ca. Steroidedax gorgoniicola* genes were screened against the NCBI nr v4 database (downloaded May 3^rd^ 2024) using blastp and an E-value cutoff of 0.05. Genes with more than one match to a gene from a member of Endozoicomonadaceae in the top ten hits were considered candidates of horizontal gene transfer (HGT). To test for potential HGT, we tested whether these candidate genes were closer to transposases than non-candidate genes. The distance of each gene to the nearest transposase was calculated in base pairs if they were on the same contig as a transposase. All transposases were excluded. A Wilcoxon rank sum test with continuity correction was used to test whether the HGT candidate genes’ distances to transposases were less than the rest of the genes in the genome. Further, we also tested whether HGT candidate genes were more clustered together than non-candidate genes. First, the minimum distance from each HGT candidate gene to the nearest HGT candidate gene was measured if another HGT candidate gene was identified on the same contig. This was repeated 10,000 times using a randomly selected set of non-candidate genes of the same magnitude as the HGT gene set. A p-value was calculated as the proportion of iterations in which the median minimum distance between genes was less than that of the HGT candidate gene set. Resolved gene trees from orthofinder were manually examined for structure indicative of HGT.

## Acknowledgements

We would like to thank the crews of all research vessels, ROV teams, and the other scientists involved with sampling.

## Data Availability

The raw sequence data and assemblies for *Ca. Steroidedax gorgoniicola* and *Ca. Nereiplasma swiftiae* are available on the NCBI database under BioProject number PRJNA875098. Amino acid sequences of predicted genes and the resolved gene trees of HGT candidate genes can be found on Figshare under these doi’s: 10.6084/m9.figshare.33170705, 10.6084/m9.figshare.33170699, 10.6084/m9.figshare.33170711.

