## Supplemental Figures for "*Ca. Steroidedax gorgoniicola,* a heterotrophic coral associate with horizontally acquired genes from Endozoicomonadaceae"

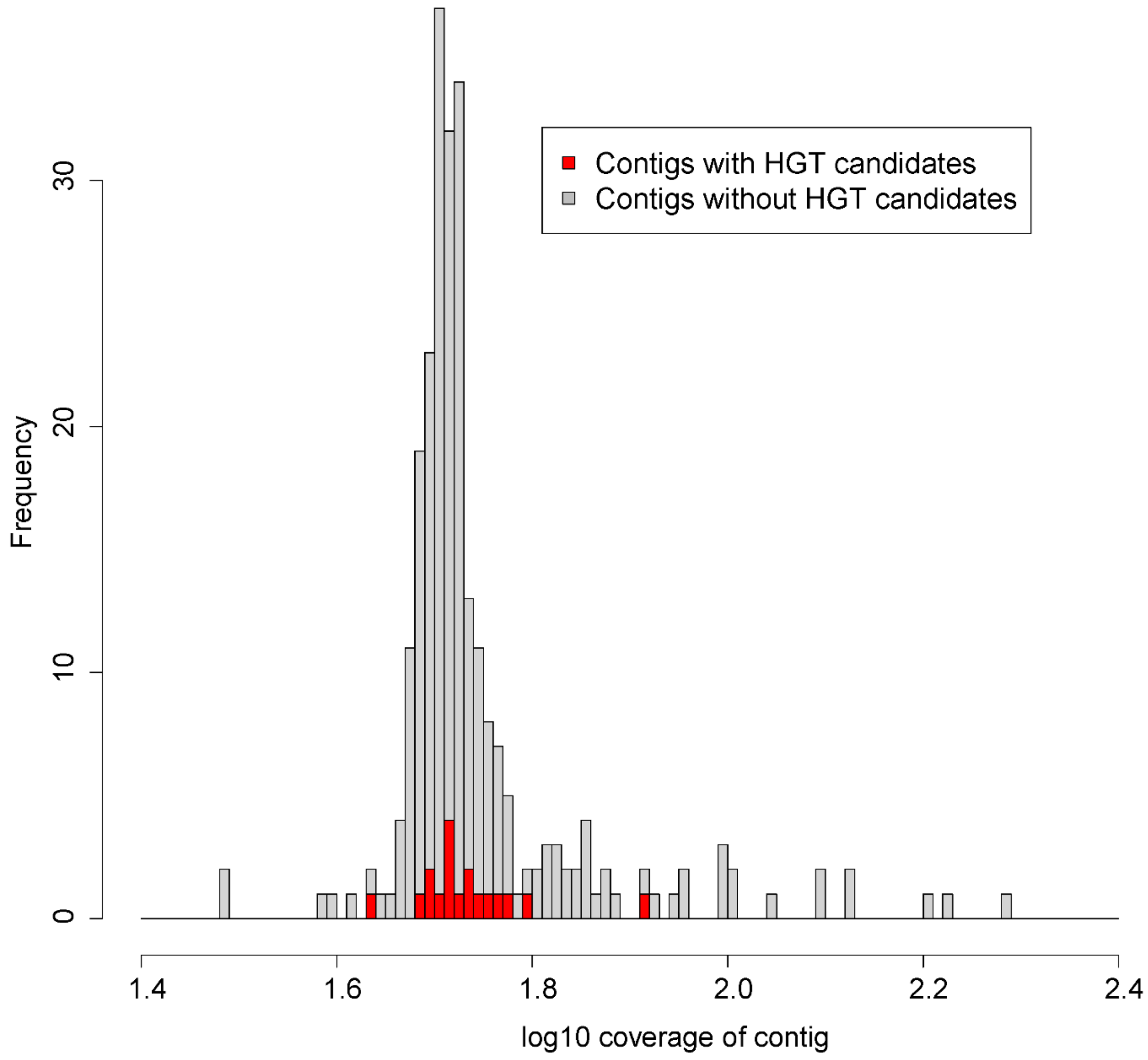


*Figure S1*

Histogram of the depth of coverage of all contigs colored depending on whether the contig contained an HGT candidate gene or not.

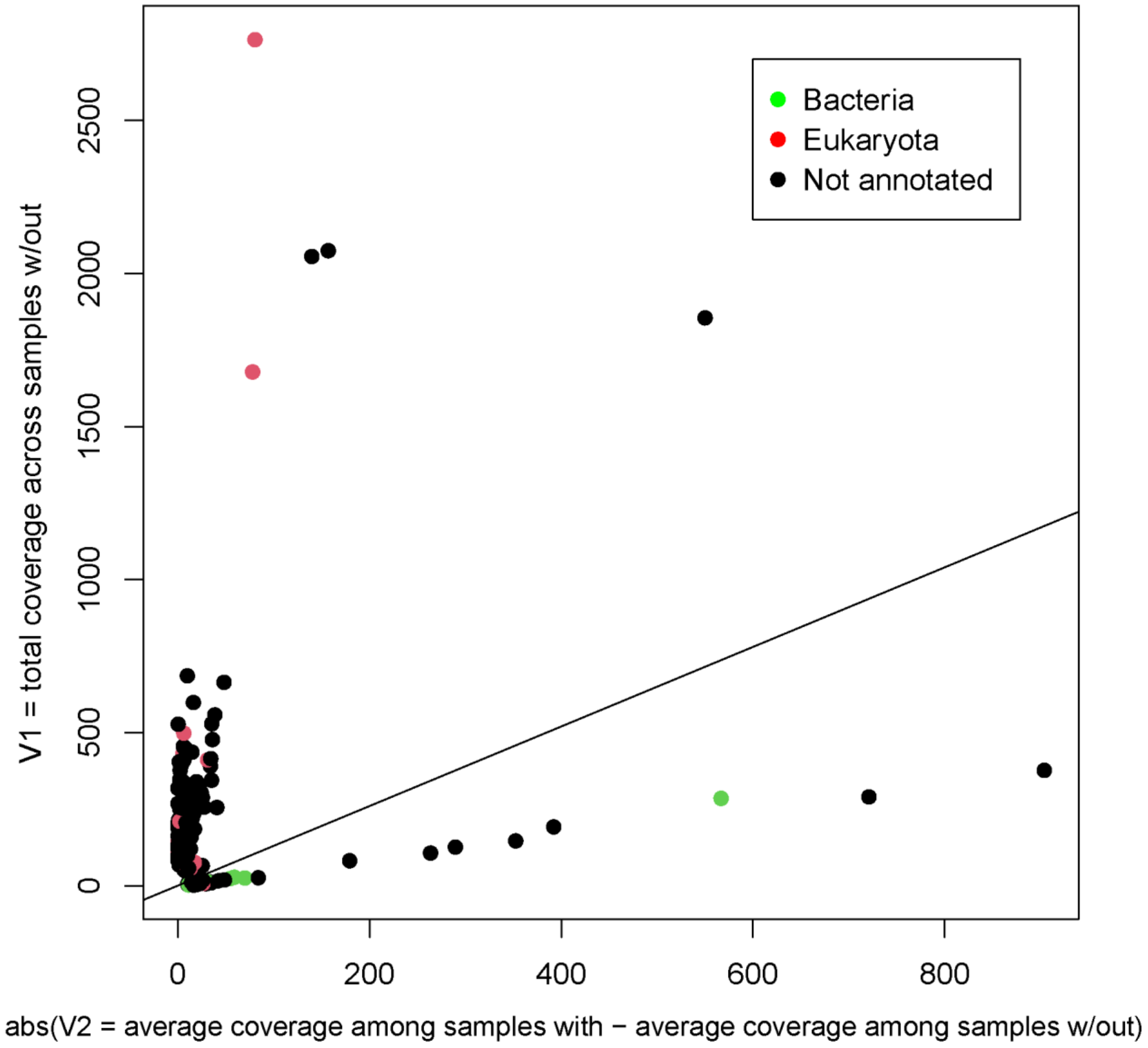


*Figure S2*

V1 versus V2 for all contigs colored by blobology classifications with a line denoting where V1/abs(V2) < 1.3. V1 is the sum of the depths of coverage of a contig across the five samples without BD1-7 and V2 is the average depth of coverage of that contig among the 4 samples with BD1-7 minus the average among the 5 samples without BD1-7

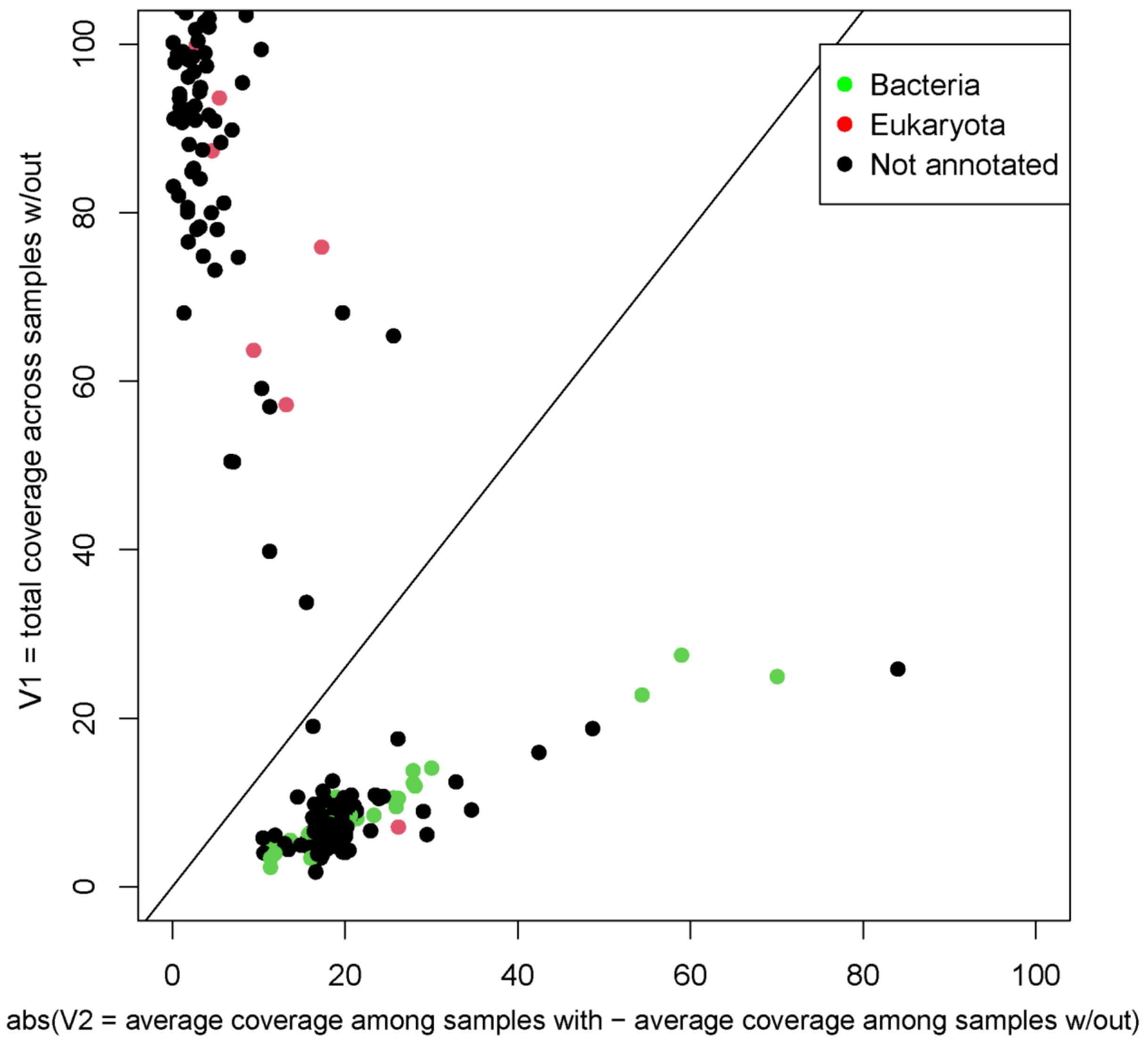


*Figure S3*

A zoomed in view of V1 versus V2 for all contigs colored by blobology classifications with a line denoting where V1/abs(V2) < 1.3. V1 is the sum of the depths of coverage of a contig across the five samples without BD1-7 and V2 is the average depth of coverage of that contig among the 4 samples with BD1-7 minus the average among the 5 samples without BD1-7
